# Towards Ecological Measurement of Complex Cognitive Processes: Functional-Near Infrared Spectroscopy of Brain Activity During Reading

**DOI:** 10.1101/2023.12.13.571603

**Authors:** Marta Čeko, Leanne Hirshfield, Emily Doherty, Rosy Southwell, Sidney D’Mello

## Abstract

Functional magnetic resonance imaging (fMRI) has provided unparalleled insights into the fundamental neural mechanisms governing human cognition, including complex processes such as reading. Here, we leverage the wealth of prior fMRI work to capture reading outside the MRI scanner using functional near infra-red spectroscopy (fNIRS). In a large sample of participants (n = 82) we observe significant prefrontal and temporal fNIRS activations during reading, which are largely reliable across participants, therefore providing a robust validation of prior fMRI work on reading-related language processing. These results lay the groundwork towards developing adaptive systems capable of assisting these higher-level processes, for example to support readers and language learners. This work also contributes to bridging the gap between laboratory findings and real-world applications in the realm of cognitive neuroscience.

## Introduction

Over the last three decades, functional magnetic resonance imaging (fMRI) has offered unprecedented insights into the fundamental neural mechanisms underlying human cognition, including complex high-level cognitive processes. Reading provides a unique context to investigate brain activity during complex processing because it involves a carefully choreographed, dynamic interaction between the reader, the text, and context ^1^. Specifically, reading entails translating a series of abstract symbols (i.e., letters) into associated units (words), extracting meanings from them (semantics), and integrating those meanings into a coherent representation of text, often going beyond the text by incorporating prior knowledge, inferences, and predictions ^2,3^. This involves several mental operations, such as parsing letter forms into words, interpreting the strings of words (i.e., sentences) based on semantic context and syntax ^4–6^, establishing connections within the text (e.g., connecting a pronoun to a previously seen noun^7^), and linking textual representations to information stored in long-term-memory (e.g., retrieving a memory associated with an event discussed in the text ^8^).

Even though reading relies on a diverse set of computations implemented across distributed brain systems, including those involved in domain-general mental processes (i.e., cognitive control, perception, memory), fMRI studies using group-level statistics have revealed consistent engagement of specific brain regions forming a distinct ‘language network’ that appears to be selective for core high-level linguistic processes compared to more domain-general processes ^9–24^. Nevertheless, the boundaries of this functional language network in the brain are blurred^17,25^, partly due to the highly distributed nature of activations covering adjacent brain regions and partly because reading is a highly individualized process^1^. Readers might engage different processes and consequently different brain regions, and these might deviate from the functional language network identified using traditional group level statistics ^17,25^. Thus, considering individual (reader-level) variation in neural recruitment and the level of correspondence with the language network identified in group- or population-level studies might reveal greater specificity per individual, which is critical to the investigation of brain mechanisms supporting reading and language processing ^9,10,16,17,26^.

To this end, Fedorenko et al. ^17^ developed a language localizer task designed to robustly define (‘localize’) language-sensitive functional brain regions (using fMRI) pertaining to different aspects of reading (e.g., identifying words, parsing sentences) distinct from those supporting non-linguistic processes. This work focused on individual participants with particular consideration given to the level of individual overlap with the overall functional language network using group-constrained participant-specific analyses ^17,27^. Subsequent studies have found these regions to be activated across the majority of participants engaged in high-level linguistic processing, to be replicable within participants over time and to have overall correspondence across participants ^16,17^. The functional language network also is robust to variations in language stimuli and presentation (e.g., visual, audio), and is consistent across many of the world’s spoken languages ^25,28^.

This body of research builds the necessary foundation for investigating reading processes in the brain. However, this work has been conducted in the fMRI scanner, which is not suited for measurement of brain function in more naturalistic settings which extends beyond reading single words or sentences into the realm of connected texts and multiple document comprehension ^29^. Therefore, fMRI also has limited utility as a real-time measurement system that could be integrated with adaptive technology to facilitate reading in real-world scenarios.

Conversely, functional near-infrared spectroscopy (fNIRS) has recently emerged as a non-invasive brain measurement technique that holds great potential for measurement of higher-level cognitive processes in the human brain in naturalistic settings^30–33^. Yet, to the best of our knowledge, it has yet to be validated as a reliable measure of the underlying neural processes implicated in reading comprehension. As a step in this direction, we use fNIRS to record brain activity during reading in participants performing the language localizer task. We show in a large sample of 82 healthy young adults that we can robustly capture cortical higher-level reading-related processes, replicating prior work with fMRI, and extending prior fNIRS work has largely focused on lower-level reading processes (i.e., reading single words) in small participant samples ^34–39^. In doing so, we validate fNIRS as a viable approach to measure complex brain processes implicated in reading comprehension outside the scanner, opening the door for more basic research and real-time personalized interventions to enhance reading outcomes (e.g., ^40^).

## Methods

### Participants

Ninety-five young adults participated in this study. The institutional review board of the University of Colorado Boulder approved the study, and all participants provided written consent. Twelve participants were excluded from data analysis due to technical issues with fNIRS data collection (unrecoverable missing trigger values), for a final analyzed sample of 82 participants (age mean (SD) 22.1 years (4.6); 28 male, 52 female, 2 other; 63 White, 9 Asian, 6 Hispanic, 4 Other).

### Language Localizer Task

The task was based on Fedorenko’s original localizer task using visual text stimuli to localize language processing in the brain ^17^. The task comprised four conditions manipulating word-level and sentence-level processing (Fig. 1a), presented in a block design of 1 block per condition, 20 text stimuli per block separated by 10s, in randomized order. Each text stimulus consisted of 12 words successively presented at fixation for 156ms, with 2 sec between sentences.

**Figure 1.**
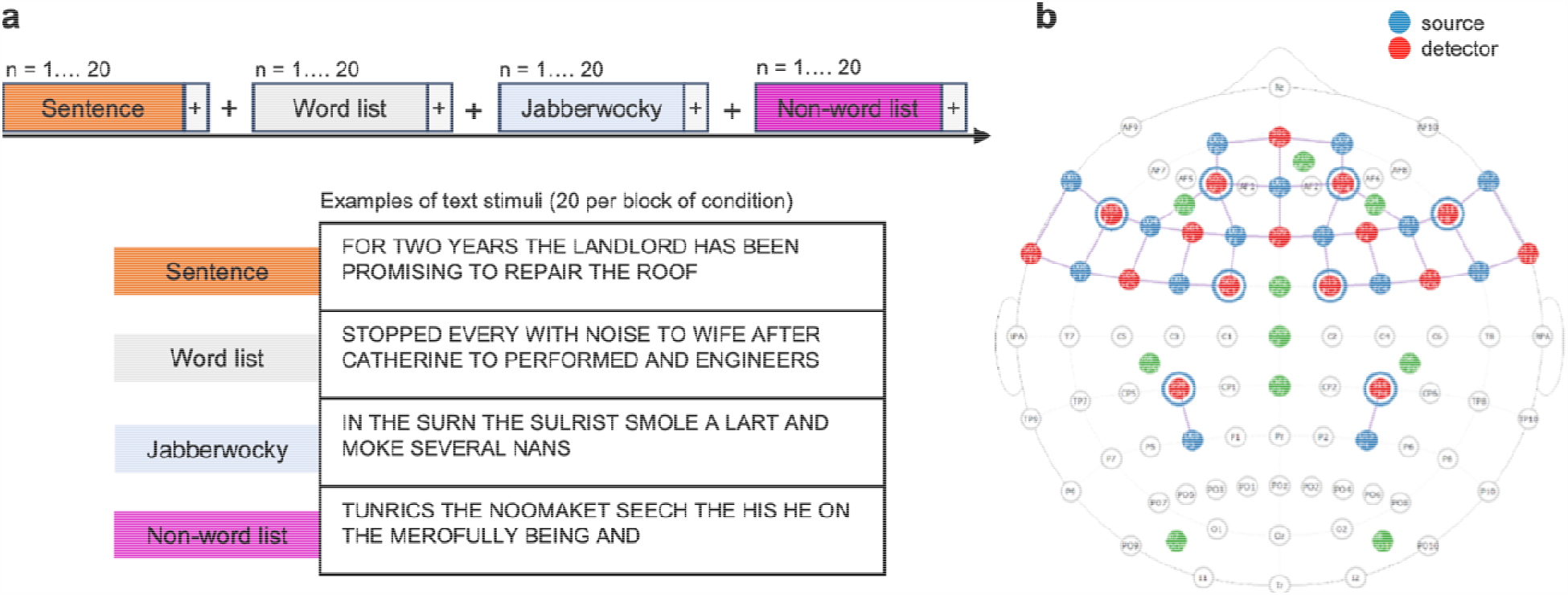
Study design and fNIRS montage. a) Experimental design for the language localizer task. Each participant was presented with 4 types of conditions (sentence, word list, jabberwocky, non-word lists), presented in a random order. Each condition consisted of 20 text stimuli, with examples shown in the figure. b) The 42-channel fNIRS montage. Each channel consisted of a source-detector pair, represented by a colored line. Note: Green circles represent EEG channels, not analyzed in this study.

The four conditions included **Sentence, Word list** (generated by shuffling words in sentences), **Jabberwocky** sentences (sentences where all the content words were replaced by pronounceable nonwords, like, “sulrist”) and **Non-word list** (shuffled jabberwocky sentences). The **Sentence** condition involves accessing the meaning of individual words (lexical semantic processing), relating the words to one another (syntactic processing), and then deriving the meaning of the entire sentence (sentence-level semantic processing). The **Word list** condition engages semantic lexical processing, but not syntactic processing, because individual lexical items (words) cannot be combined into more complex representations to form a sentence. The **Jabberwocky** condition engages syntactic processing as it follows the rules of English syntax to form a sentence, but not semantic lexical processing, as it includes nonsense words. The **Non-word list** condition contains a nonsense string of pronounceable non-words and therefore does not engage syntactic nor sentence-level semantic processing.

The comparison between sentences and nonword lists allows identification of regions in the brain that are sensitive to word- and sentence-level meaning and sentence structure and has thus been used in the fMRI language literature to capture higher-level linguistic processes, such as those that underlie reading of texts. Our contrast of interest (and the main contrast investigated in the original localizer study ^17^), is [**Sentence – Non-word list**]. This contrsast targets regions in the brain that are sensitive to word- and sentence-level meaning and sentence structure. This contrast has thus been used in the fMRI studies to capture higher-level linguistic processes, such as those that underlie reading of texts, as these involve retrieving the meanings of individual words and combining these lexical meanings into more complex structural representations (sentences and texts).

### fNIRS data acquisition

We collected fNIRS data on 95 participants presented with the language localizer task using the channel montage in Fig. 1b as part of a larger study (not analyzed here). Data collection was conducted using the NirSport 16 x 16 system (NIRx, Berlin, Germany) with a sampling rate of 10.2 Hz.

The fNIRS montage was designed to target brain areas which have been found to play a role in reading comprehension and language processing. The channel montage was guided by the functional language network, defined initially using the language localizer task by Fedorenko et al. ^10,17^ and refined in other studies ^11,12,15,16,22,23,28,41^, and more generally by meta-analytic functional brain maps associated with the terms ‘language’, ‘reading’ and ‘reading comprehension’ derived from Neurosynth (neurosynth.org), as well as regions associated with attention, integration and executive functioning. Specifically, the montage of 42 channels spanned prefrontal, parietal and temporal cortices, including bilaterally the inferior frontal gyrus (IFG), and orbital IFG (IFGorb), middle frontal gyrus (MFG), superior frontal gyrus (SFG), angular gyrus (AngG), and anterior and posterior temporal gyri (AntTemp, PostTemp).

### fNIRS data processing

All fNIRS preprocessing was conducted in the NIRS Brain AnalyzIR Toolbox (version 837) ^42^ in MATLAB (version 2021a). Briefly, the raw voltage data were resampled to 5.1Hz, converted to optical density, and then converted to changes in oxygenated (HbO) and deoxygenated (HbR) hemoglobin using the modified Beer-Lambert Law^43^, followed by a bandpass filter (.01-.5 Hz) to remove systemic artifacts including heart rate, respiration, and blood pressure. HbO and HbR timeseries data were used to calculate subject-level (first-level) statistics using a General Linear Model (GLM) with an autoregressive pre-whitening filter to account for serially correlated errors and downweigh outliers due to motion artifacts^44^. The first level GLM thus estimated the [sentences – nonwords] contrast per participant averaged across 20 text stimuli.

### Group-level analysis

The resulting first-level statistic contains the subject-level regression coefficients (beta values for the contrast) as well as their corresponding error-covariance matrices. Using the output from the first-level analysis, a second-level (group-level) model was calculated, using the full covariance from the first-level models, to perform a weighted least-squares regression and generate group level statistics on the [sentences – nonwords]. To control for multiple comparisons, false discovery rate (FDR) correction was used with the significance level set at 0.05 (q ≤ 0.05) ^45^. Analysis results are t-values resulting from the contrasts.

### Individual variability and correspondence with fMRI-derived functional language localizer

Because individual differences can attenuate the effects of localizers in group-level statistics ^17^, we next computed for each fNIRS channel the numer of participants with a significant activation in this channel. We defined the top 10% channels as regions high between-subject reliability. The bottom 10% were defined as regions showing low between-subject reliability. This served to test whether the [sentence-nonword] contrast reliably engages the same regions across participants.

To assess the degree of spatial overlap with the functional language network we mapped the top 10% and bottom 10% channels onto the current functional language localizer map downloaded from evlab.mit.edu//funcloc/ (and encompassing bilaterally the IFG, IFGorb, MFG, AntTemp, MidTemp and PostTemp, but not the SFG included in the original localizer ^17^). The MNI coordinates corresponging to the fNIRS channels were overlaid onto the localizer map on top of a standard anatomical MRI template in MNI space. We expected the fNIRS channels with high between-subject reliability to have high anatomical overlap with the localizer. As a control, we expected the bottom 10% (low inter-subject reliability) channels to show weaker overlap with the localizer.

## Results

### Group Results

Group results are depicted in Fig. 2. HbO activations have positive t-values (red in Fig. 2a) while HbR activations have negative t-values (blue in Fig. 2b). Table 1 lists these group results, with only significant activations of HbO and HbR included.

**Figure 2.**
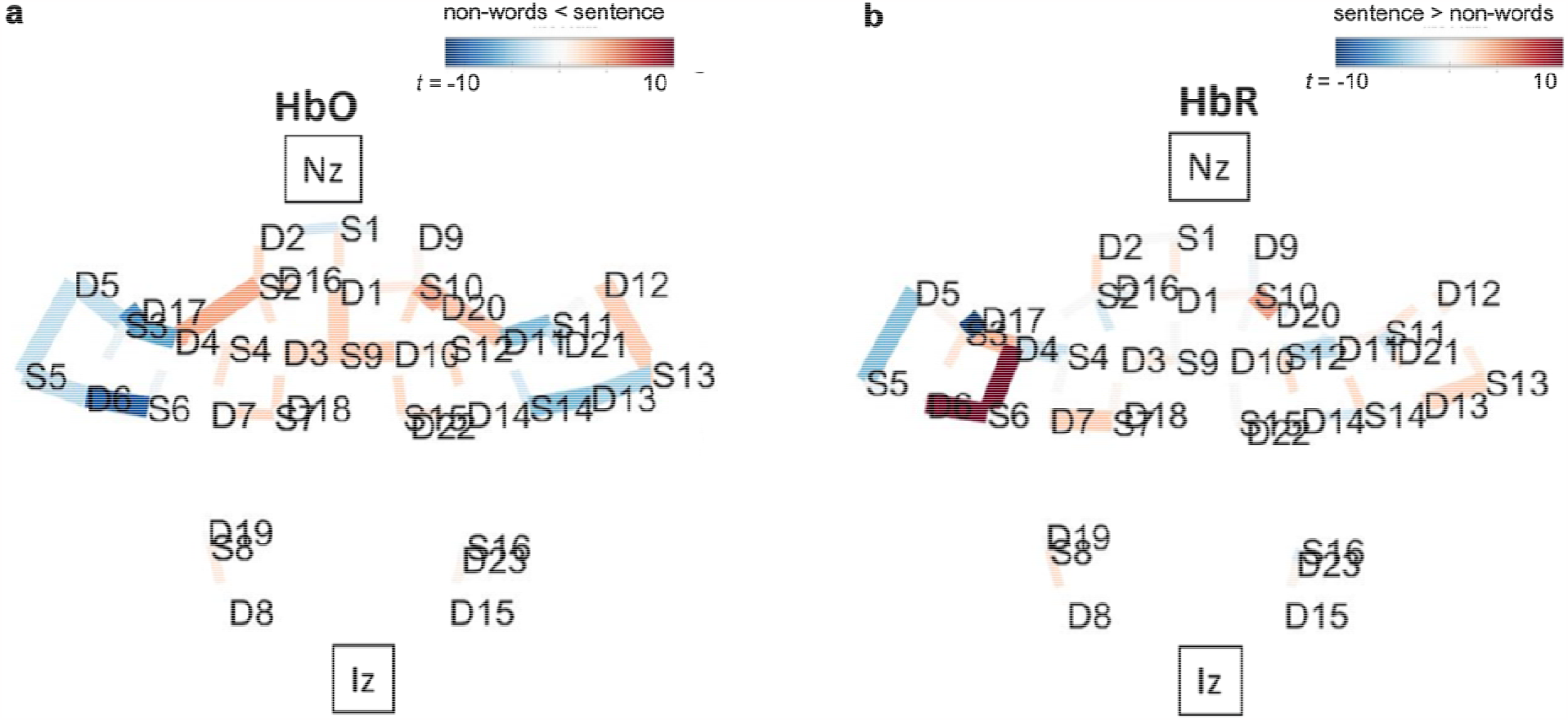
fNIRS group results for the [sentence-nonwords] contrast. Both directions of the contrast (sentences – nonwords) and its inverse (nonwords – sentences) are overlaid over a brain, with nasion (Nz) and inion (Iz) locations added for reference. Only significant channels (*q* < 0.05) are shown. **a)** For HbO, positive *t*-values (red) correspond to relatively larger activity for the first term in the contrast, and negative *t*-values (blue) correspond to larger activity for the second term. **b)** For HbR contrasts, negative *t*-values (blue) correspond to larger activation in that region.

**Table 1.** fNIRS group results for the [sentence-nonwords] contrast.

| s | d | brain region | FLL | L / R | beta (se) | t | p | q |
| --- | --- | --- | --- | --- | --- | --- | --- | --- |
| *Significant HbO activations* |  |  |  |  |  |  |  |  |
| 2 | 4 | BA 45 – IFG, Broca | IFGorb | L | 4.61 (1.12) | 4.11 | 0.00 | 0.00 |
| 9 | 1 | BA 09 – MFG<br>BA 10 – frontal pole | SFG* | mid | 7.47 (2.88) | 2.59 | 0.01 | 0.03 |
| 9 | 3 | BA 09 – MFG<br>BA 08 – SFG | SFG* | L | 5.92 (2.51) | 2.36 | 0.02 | 0.05 |
| 9 | 10 | BA 09 – MFG<br>BA 08 – SFG | SFG* | R | 7.44 (2.56) | 2.90 | 0.00 | 0.01 |
| 11 | 11 | BA 45 – IFG, Broca | IFGorb | R | 4.46 (1.35) | 3.29 | 0.00 | 0.00 |
| 13 | 12 | BA 38 – AntTemp | AntTemp | R | 5.32 (1.91) | 2.79 | 0.01 | 0.02 |
| *Significant HbR activations* |  |  |  |  |  |  |  |  |
| 4 | 4 | BA 45 – IFG, Broca<br>BA 46 – MFG | IFG | L | -2.46 (1.02) | -2.40 | 0.02 | 0.04 |
| 5 | 5 | BA 38 – AntTemp | AntTemp | L | -4.86 (1.09) | -4.47 | 0.00 | 0.00 |
| 12 | 11 | BA 45 – IFG, Broca<br>BA 46 – MFG | IFG | R | -0.85 (0.27) | -3.14 | 0.00 | 0.01 |

Notes: Only significant activations are included (significant HbO positive t-value and significant HbR negative t-values). All regions mapped onto the original group-level functional language localizer (FLL; [sentences > nonword lists]<sup>17</sup>). \* SFG was included in the original localizer but is not included in the current version of the localizer. Dfe = 5120.29. s, source; d = detector; BA, Brodman area; L / R, left / right hemisphere; se, standard error; t, t-value; p, p-value; q, FDR-corrected p-value; I / M / SFG, inferior / middle / superior frontal gyrus; (a) ITG, inferior temporal gyrus. AntTemp = anterior temporal gyrus (temporal pole).

These group level results align well with the group-level functional language network, defined initially using the language localizer task by Fedorenko et al. ^17^. Specifically, all 9 significant group-level results mapped onto Fedorenko’s language localizer parcellation, with four significant fNIRS activations (i.e., channels) mapping onto the IFG bilaterally, three onto MFG bilaterally and two onto the anterior temporal gyrus (AntTemp, bilaterally. Fedorenko’s group level contrast [sentences > nonwords list] resulted in activations in the left IFG, left IFGorb, left MFG, left SFG, bilateral AntTemp, bilateral Mid- and PostTemp, left AngG and to a lesser extent in the cerebellum ^17^.

### Individual results and correspondence with fMRI-derived functional language localizer

We also generated our statistical results per individual, because individual differences can attenuate the effects of localizers in group level statistics ^17,27^. we generated statistical results per participant on the main contrast ([sentences – nonwords]). The main contrast resulted in 681 total channels across 77 participants showing significant (q<.05) activation (either positive HbO t-values or negative HbR t-values) across all subjects. On average this resulted in just over 8 significant channels per participant (mean (SD), 8.8 (6.9), range: 1 – 39 channels).

Channels with the highest between-subject reliability (defined as the top 10%) all mapped onto inferior frontal gyri (including the left-lateralized Broca’s area; Fig. 3a, red spheres 1-4; right BA38/45, left BA45, right BA48/6, left BA38/45). Across the four channels showing the highest between-subject reliability, 55 participants (77%) had significant activation in at least one channel (Fig. 3b). These channels showed high anatomical overlap with bilateral inferior frontal areas in Fedorenko’s language localizer map. Comparable inter-subject reliability of channels was observed when split by signal type (HbO, HbR), with channels mapping to the same or similar anatomical location as the combined-signal channels in Fig. 3. Of note, next four highest-reliability channels (top 20% percent; bars 5-8 in Fig. 3a) also mapped onto IFG, IFGorb, and additionally to bilateral AntTemp. The control channels with low inter-subject reliability (defined as the bottom 10%) showed less anatomical overlap with Fedorenko’s language localizer or showed overlap with brain regions less reliably activated in language studies. Specifically, the channel mapped to SFG (blue sphere 38 in Fig. 3a) showed high overlap with the SFG, an area that was included in the initial^17^, but not in subsequent versions of the localizer (e.g. ^16,28^) and that has shown lower group-level activation in the original localizer experiments^17^, and low reliability in replication and extension studies with new participants and new language comprehension paradigms ^9,10,16,17,25,26^. Of note, the AngG (blue sphere 37 in Fig.3a) also typically shows less activation than frontal and temporal areas in studies of language processing^10,11,17,28^.

**Figure 3.**
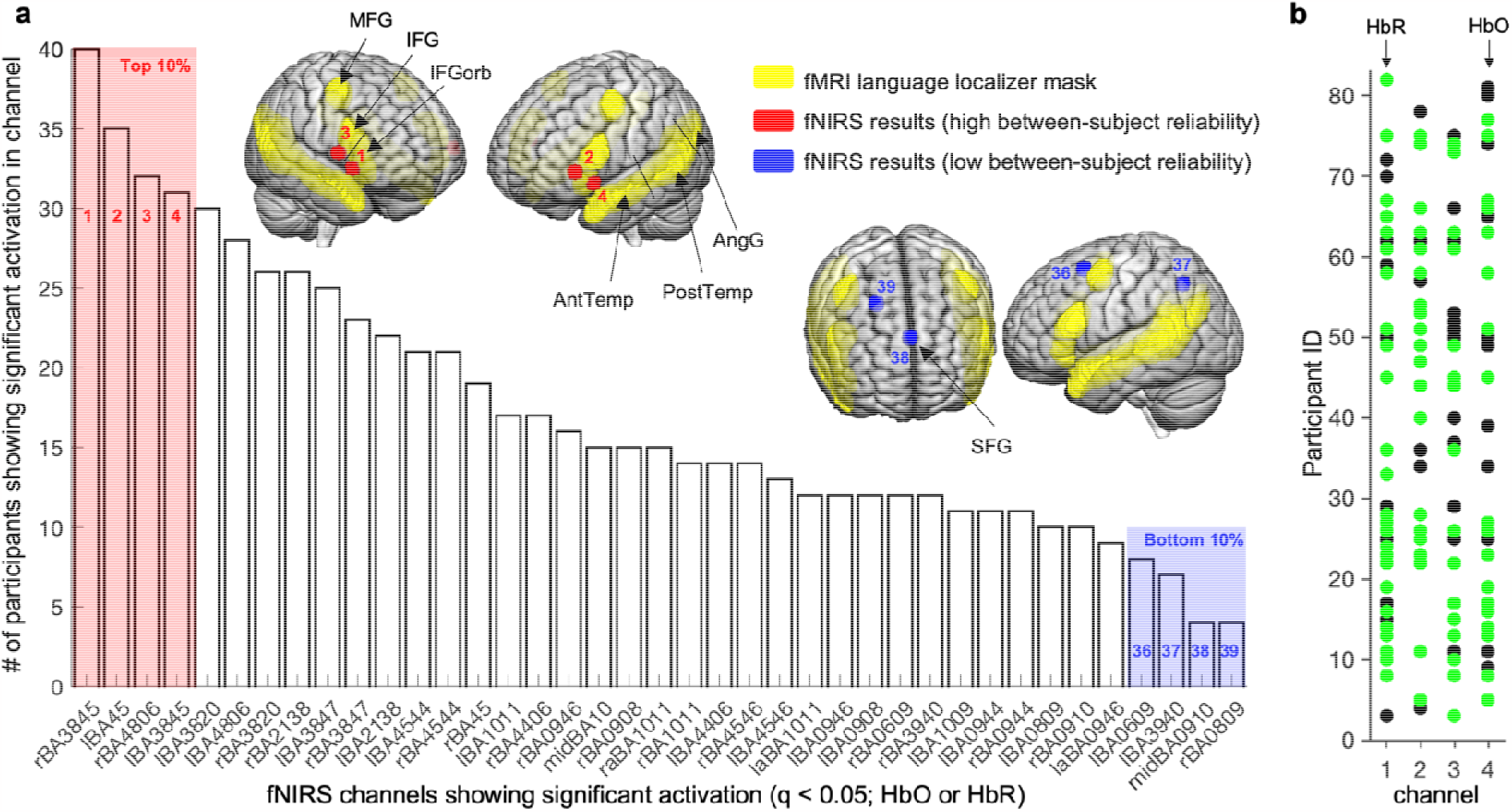
Between-subject variability in significant fNIRS activations. A) fNIRS channels - labelled with corresponding Brodmann anatomical areas - are shown in the chart with the number of participants with significant activation in that channel. The fMRI-based functional language localizer map (downloaded from evlab.mit.edu//funcloc/) is shown in yellow on the cortical surface of a standard MRI template in MNI space. The top and bottom 10% are highlighted in red and blue respectively in the chart and on the brain map. Regions (black arrows) are named using the convention used in ^17^. B) Significant channels per participant ID are displayed for the top 10% channels (HbR channels in green, HbO channels in black). MFG, middle frontal gyrus; IFG, inferior frontal gyrus; IFGorb, orbital IFG; Ant / Post Temp, anterior / posterior temporal gyrus; AngG, angular gyrus.

## Discussion

We administered an established language localizer task ^17^ with individuals equipped with fNIRS sensors to extend prior fMRI work on localizing language processing in the human brain. Our results, obtained outside the fMRI scanner, demonstrate the convergent validity of fNIRS in identifying cortical brain regions activated during linguistic processes involved in reading and underscore the potential of fNIRS for real-world measurement of reading processes. This is the largest study to date to use fNIRS to capture reading-related cortical activation, and the only one using comprehensive brain coverage, therefore providing a robust validation of prior fMRI work on reading-related language processing. Group-level and participant-level analyses validate the utility of fNIRS for capturing reading processes with the ultimate goal of tracking and enhancing reading comprehension in an individualized manner.

Our group-level results aligned with the functional brain network defined using the language localizer task ^17^ by showing activations in the inferior and mid prefrontal cortices and the anterior temporal lobe. At the participant-level, significant fNIRS activations common to most participants (i.e., having high between-subject reliability) mapped bilaterally onto the inferior prefrontal cortices and - to lesser extent - onto the bilateral anterior temporal cortices. These results are thus in line with contemporary fMRI studies of language processing which show most reliable and robust activations in inferior prefrontal and temporal regions (especially in the left hemisphere) ^10,12,13,16,17,22,23,25,41,46,47^. The fNIRS activcations showing the most variability across participants (i.e., low inter-subject reliability) also showed less overlap with the language network, by either having less anatomical overlap with the established functional localizer or showing anatomical overlap with superior frontal and posterir temporal brain areas less reliably activated in language studies ^9,10,16,17,25,26^.

The significance of this work lies in showcasing our ability to measure neural correlates of language-localizing tasks outside the traditional fMRI scanner. This has implications for the domains of education and human-computer interaction, and applications such as usability testing of reading comprehension assessments, design of adaptive learning systems, and medical diagnostics related to reading. By leveraging foundational fMRI studies, our research takes a crucial step towards developing adaptive systems capable of assisting and enhancing reading outcomes, thereby bridging the gap between laboratory findings and real-world applications in cognitive neuroscience. Moroever, the finding that the majority of participants (71%) showed activation in the inferior frontal cortex, suggests that wearable, low-cost versions of fNIRS – with a few channels concentrated onto the prefrontal cortex - might be sufficient to capture brain correlates of reading in ecological settings. This finding also provides support to ongoing efforts to develop cost-effective ultra-wearable and ultra-unobtrusive fNIRS devices to support human functioning across domains.

This work has several limitations. First, while we followed standard procedures for removing motion and physiological artifacts from the fNIRS signal, we did not additionally regress out short-channel signal –which might have improved brain activity estimates by removing systemic noise from the signal of interest ^48^. Second, the population used in this study (a homogenous sample of young, predominantly white, adults - mostly undergraduate students) is a limitation towards the generalizability of our findings.

Future work can build on these findings to use fNIRS to measure complex reading-related and other cognitive processes in real-time, including mind wandering ^49^, inferencing ^8^, and error correction (i.e. regressions) ^50^. This can be followed by using more naturalistic reading paradigms such as reading of long connected texts^51^ with the goals of learning and retention, rather than individual sentences that the bulk of fMRI work on language processing is based on. There is also a rich literature base on eye movements and reading (e.g., ^51,52^), which can be complemented with real-time fNIRS measurement to help disambiguate particular eye movements (e.g., ^21^). A longer-term goal is to use multimodal measurement of eye movements and fNIRS to select real-time intelligent adaptations to support individuals in personalized ways during reading. By demonstrating for the first time that higher-order processes implicated in reading can be reliably measured outside the scanner, we open the door to new research opportunities and application areas.

## Acknowledgements

We thank Trevor Grant for assistance in the early stages of the project. This research was supported by the National Science Foundation (DRL 1920510). The opinions expressed are those of the authors and do not represent views of the funding agency.

## Author contributions

Conceptualization: S.D.,L.H., data collection: E.D., R.S., analysis and interpretation. M.C., L.H., E.D., R.S., writing - original draft, M.C., L.H..; writing - review & editing, M.C., L.H., E.D., R.S., S.D.

## Data and code availability statement

Source data and code needed to reproduce the results presented in Figs. 2-3 and Table 1 will be available at https://github.com/emotive-computing/EML-localizer upon publication.

## References

1. Snow, C. Reading for Understanding: Toward an R&D Program in Reading Comprehension. RAND Corporation (2002).

2. Kintsch, W. The role of knowledge in discourse comprehension: a construction-integration model. Psychol. Rev. 95, 163–182 (1988).

3. Mcnamara, D. S. & Magliano, J. Toward a comprehensive model of comprehension. Psychology of learning and motivation.

4. Jurafsky, D. A probabilistic model of lexical and syntactic access and disambiguation. Cognitive Science 20, 137–194 (1996).

5. Spivey, M. J. & Tanenhaus, M. K. Syntactic ambiguity resolution in discourse: Modeling the effects of referential context and lexical frequency. Journal of Experimental Psychology: Learning, Memory, and Cognition 24, 1521 (1998).

6. Tanenhaus, M. K. & Trueswell, J. C. Sentence comprehension. in Speech, Language, and Communication (eds. Miller, J. L. & Eimas, P. D.) (Academic Press, 1995).

7. Dell, G. S., McKoon, G. & Ratcliff, R. The activation of antecedent information during the processing of anaphoric reference in reading. Journal of Memory and Language 22, 121 (1983).

8. McNamara, D. S. If Integration Is the Keystone of Comprehension: Inferencing Is the Key. Discourse Processes 56, 86–91 (2021).

9. Fedorenko, E. & Thompson-Schill, S. L. Reworking the language network. Trends in cognitive sciences 18, 120–126 (2014).

10. Fedorenko, E., Behr, M. K. & Kanwisher, N. Functional specificity for high-level linguistic processing in the human brain. Proceedings of the National Academy of Sciences 108, 16428–16433 (2011).

11. Mineroff, Z., Blank, I. A., Mahowald, K. & Fedorenko, E. A robust dissociation among the language, multiple demand, and default mode networks: Evidence from inter-region correlations in effect size. Neuropsychologia 119, 501–511 (2018).

12. Ferstl, E. C., Neumann, J., Bogler, C. & von Cramon, D. Y. The extended language network: A meta-analysis of neuroimaging studies on text comprehension. Human Brain Mapping 29, 581–593 (2008).

13. Diachek, E., Blank, I., Siegelman, M., Affourtit, J. & Fedorenko, E. The domain-general multiple demand (MD) network does not support core aspects of language comprehension: A large-scale fMRI investigation. J. Neurosci. 40, 4536–4550 (2020).

14. Schrimpf, M. et al. The neural architecture of language: Integrative modeling converges on predictive processing. Proc. Natl. Acad. Sci. U. S. A. 118, e2105646118 (2021).

15. Blank, I., Kanwisher, N. & Fedorenko, E. A functional dissociation between language and multiple-demand systems revealed in patterns of BOLD signal fluctuations. J. Neurophysiol. 112, 1105–1118 (2014).

16. Mahowald, K. & Fedorenko, E. Reliable individual-level neural markers of high-level language processing: A necessary precursor for relating neural variability to behavioral and genetic variability. Neuroimage 139, 74–93 (2016).

17. Fedorenko, E., Hsieh, P.-J., Nieto-Castañón, A., Whitfield-Gabrieli, S. & Kanwisher, N. New method for fMRI investigations of language: defining ROIs functionally in individual subjects. Journal of neurophysiology 104, 1177–1194 (2010).

18. DeWitt, I. & Rauschecker, J. P. Wernicke’s area revisited: Parallel streams and word processing. Brain Lang. 127, 181–191 (2013).

19. Binder, J. R. Current Controversies on Wernicke’s Area and its Role in Language. Curr. Neurol. Neurosci. Rep. 17, (2017).

20. Pallier, C., Devauchelle, A.-D. & Dehaene, S. Cortical representation of the constituent structure of sentences. Proc. Natl. Acad. Sci. U. S. A. 108, 2522–2527 (2011).

21. Henderson, J. M., Choi, W., Luke, S. G. & Desai, R. H. Neural correlates of fixation duration in natural reading: evidence from fixation-related fMRI. Neuroimage 119, 390–397 (2015).

22. Hsu, C.-T., Clariana, R., Schloss, B. & Li, P. Neurocognitive signatures of naturalistic reading of scientific texts: a fixation-related fMRI study. Scientific reports 9, 1–16 (2019).

23. Swett, K. et al. Comprehending expository texts: the dynamic neurobiological correlates of building a coherent text representation. Frontiers in Human Neuroscience 7, 853 (2013).

24. Moss, J. & Schunn, C. D. Comprehension through explanation as the interaction of the brain’s coherence and cognitive control networks. Front. Hum. Neurosci. 9, 562 (2015).

25. Malik-Moraleda, S. et al. An investigation across 45 languages and 12 language families reveals a universal language network. Nature Neuroscience 25, 1014–1019 (2022).

26. Braze, D. et al. Unification of sentence processing via ear and eye: An fMRI study. cortex 47, 416–431 (2011).

27. Nieto-Castañón, A. & Fedorenko, E. Subject-specific functional localizers increase sensitivity and functional resolution of multi-subject analyses. Neuroimage 63, 1646–1669 (2012).

28. Scott, T. L., Gallée, J. & Fedorenko, E. A new fun and robust version of an fMRI localizer for the frontotemporal language system. Cogn. Neurosci. 8, 167–176 (2017).

29. Richter, T. & Maier, J. Comprehension of multiple documents with conflicting information: A two-step model of validation. Educ. Psychol. 52, 148–166 (2017).

30. Doherty, E. J. et al. Interdisciplinary views of fNIRS: Current advancements, equity challenges, and an agenda for future needs of a diverse fNIRS research community. Front. Integr. Neurosci. 17, 1059679 (2023).

31. Eloy, L., Doherty, E. J., Spencer, C. A., Bobko, P. & Hirshfield, L. Using fNIRS to identify transparency- and reliability-sensitive markers of trust across multiple timescales in collaborative human-human-agent triads. Front. Neuroergonomics 3, (2022).

32. Grant, T. et al. A neurophysiological sensor suite for real-time prediction of pilot workload in operational settings. in HCI International 2020 – Late Breaking Papers: Cognition, Learning and Games 60–77 (Springer International Publishing, 2020).

33. Hirshfield, L., Wickens, C., Doherty, E., Spencer, C., Williams, T., Hayne, L. Toward Workload-Based Adaptive Automation: The Utility of fNIRS for Measuring Load in Multiple Resources in the Brain. Preprint at (2023).

34. Hofmann, M. J. et al. Occipital and orbitofrontal hemodynamics during naturally paced reading: an fNIRS study. Neuroimage 94, 193–202 (2014).

35. Tse, C.-Y. et al. Imaging cortical dynamics of language processing with the event-related optical signal. Proc. Natl. Acad. Sci. U. S. A. 104, 17157–17162 (2007).

36. Kennan, R. P., Kim, D., Maki, A., Koizumi, H. & Constable, R. T. Non-invasive assessment of language lateralization by transcranial near infrared optical topography and functional MRI. Hum. Brain Mapp. 16, 183–189 (2002).

37. Bisconti, S., Di Sante, G., Ferrari, M. & Quaresima, V. Functional near-infrared spectroscopy reveals heterogeneous patterns of language lateralization over frontopolar cortex. Neurosci. Res. 73, 328–332 (2012).

38. Lo, Y. L. et al. Correlation of near-infrared spectroscopy and transcranial magnetic stimulation of the motor cortex in overt reading and musical tasks. Motor Control 13, 84–99 (2009).

39. Quaresima, V., Bisconti, S. & Ferrari, M. A brief review on the use of functional nearinfrared spectroscopy (fNIRS) for language imaging studies in human newborns and adults. Brain Lang. 121, 79–89 (2012).

40. Mills, C., Gregg, J., Bixler, R. & D’Mello, S. K. Eye-Mind Reader: An Intelligent Reading Interface that Promotes Long-term Comprehension by Detecting and Responding to Mind Wandering. Human-Computer Interaction 36, 306–302 (2021).

41. Yarkoni, T., Speer, N. K. & Zacks, J. M. Neural substrates of narrative comprehension and memory. Neuroimage 41, 1408–1425 (2008).

42. Santosa, H., Zhai, X., Fishburn, F. & Huppert, T. The NIRS Brain AnalyzIR Toolbox. Algorithms 11, (2018).

43. Strangman, G., Franceschini, M. A. & Boas, D. A. Factors affecting the accuracy of nearinfrared spectroscopy concentration calculations for focal changes in oxygenation parameters. NeuroImage 18, 865–879 (2003).

44. Meidenbauer, K. L., Choe, K. W., Cardenas-Iniguez, C., Huppert, T. J. & Berman, M. G. Load-Dependent Relationships between Frontal fNIRS Activity and Performance: A Data-Driven PLS Approach. bioRxiv 2020.08.21.261438 (2020).

45. Benjamini, Y. & Hochberg, Y. Controlling the False Discovery Rate: A Practical and Powerful Approach to Multiple Testing. Journal of the Royal Statistical Society: Series B (Methodological) 57, 289–300 (1995).

46. Schuster, S., Hawelka, S., Himmelstoss, N. A., Richlan, F. & Hutzler, F. The neural correlates of word position and lexical predictability during sentence reading: evidence from fixation-related fMRI. Language, Cognition and Neuroscience 1–12 (2019).

47. Yarkoni, T., Speer, N. K., Balota, D. A., McAvoy, M. P. & Zacks, J. M. Pictures of a thousand words: investigating the neural mechanisms of reading with extremely rapid eventrelated fMRI. Neuroimage 42, 973–987 (2008).

48. Wyser, D. et al. Short-channel regression in functional near-infrared spectroscopy is more effective when considering heterogeneous scalp hemodynamics. Neurophotonics 7, 035011 (2020).

49. D’Mello, S. K. & Mills, C. S. Mind wandering during reading: An interdisciplinary and integrative review of psychological, computing, and intervention research and theory. Language and Linguistics Compass 15, e12412 (2021).

50. Rayner, K., Pollatsek, A., Ashby, J. & Clifton, C., Jr. The Psychology of Reading. (Psychology Press, 2012).

51. Southwell, R., Gregg, J., Bixler, R. & D’Mello, S. K. What Eye Movements Reveal about Later Comprehension of Long, Connected Texts. Cognitive Science 44, e12905 (2020).

52. Rayner, K. Eye movements in reading: Models and data. Journal of Eye Movement Research 2, 1–10 (2009).

